# Single cell transcriptome analysis of human pancreas reveals transcriptional signatures of aging and somatic mutation patterns

**DOI:** 10.1101/108043

**Authors:** Martin Enge, H. Efsun Arda, Marco Mignardi, John Beausang, Rita Bottino, Seung K. Kim, Stephen R. Quake

## Abstract

As organisms age, cells accumulate genetic and epigenetic changes that eventually lead to impaired organ function or catastrophic transformation such as cancer. Since aging appears to be a stochastic process of increasing disorder^1^ cells in an organ will be individually affected in different ways - thus rendering bulk analyses of postmitotic adult tissues difficult to characterize. Here we directly measure the effects of aging in primary human tissue by performing single-cell transcriptome analysis of 2544 human pancreas cells from eight donors spanning six decades of life. We find that islet cells from older donors have increased levels of molecular disorder as measured both by noise in the transcriptome and by the number of cells which display inappropriate hormone expression, revealing a transcriptional instability associated with aging. By further analyzing the spectrum of somatic mutations in single cells, we found a specific age-dependent mutational signature characterized by C to A and C to G transversions. These mutations are indicators of oxidative stress and the signature is absent in single cells from human brain tissue or in a tumor cell line. We have used the single cell measurements of transcriptional noise and mutation level to identify molecular pathways correlated with these changes that could influence human disease. Our results demonstrate the feasibility of using single-cell RNA-seq data from primary cells to derive meaningful insights into the genetic processes that operate on aging human tissue and to determine molecular mechanisms coordinated with these processes.

## Main

Aging in higher-order metazoans is the result of a gradual accumulation of cellular damage, which eventually leads to a decline in tissue function and fitness ^1^. Since the fundamental processes involved in aging affect single cells in a stochastic manner, they have been difficult to study systematically in primary human tissue. Studies of selected genes in mice indicate that aging postmitotic cells of the heart display a transcriptional instability ^2^ that is not observed in actively renewing cell populations such as those of the hematopoietic system ^3^. An accumulation of genetic aberrations has been suggested to underlie transcriptional dysregulation by affecting promoter and enhancer elements as well as exonic sequences ^4^. However, due to technical constraints it has previously been difficult to study these processes in human tissue or at the whole transcriptome level.

In order to investigate the effect of physiological aging on pancreatic epithelial cells, we obtained pancreata from eight previously-healthy donors operationally defined as juvenile (ages 1 month, 5 years and 6 years), young adult (ages 21 and 22) and adult/middle aged (ages 38, 44 and 54). Single pancreatic cells were purified by flow cytometry and their mRNA expression analyzed using single-cell RNA-Seq (scRNA-seq) ^5^, with the quality of individual cells assessed using an automated quality control pipeline (see *Methods* for details). Dimensionality reduction analysis (tSNE) of data from all donors led to consistent clustering of different cell types into distinct regions (Fig. 1a), indicating good and an absence of donor or sequencing related batch effects.

**Figure 1.**
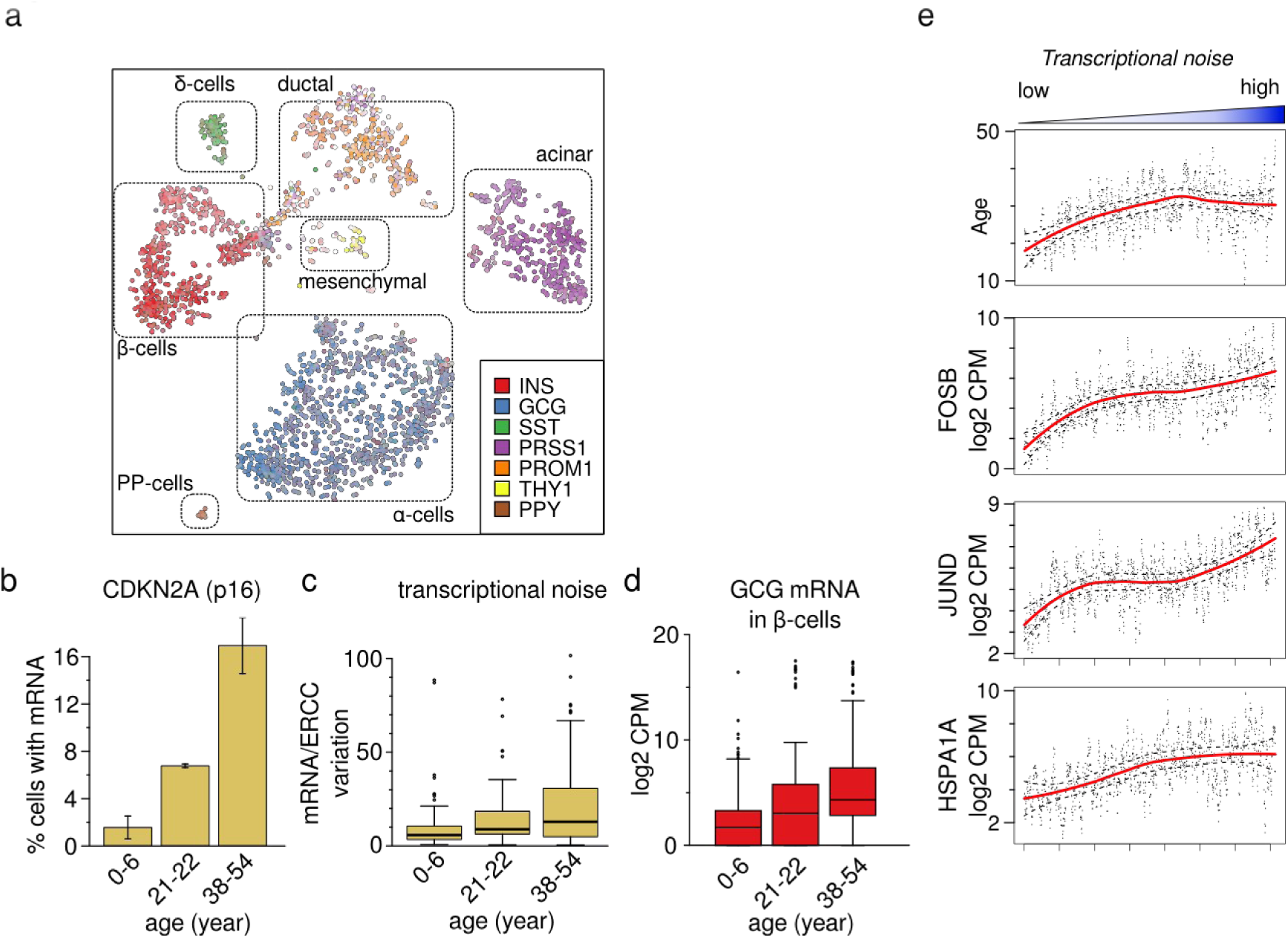
A Comprehensive survey of single cells sampled from human pancreas across different ages. a) tSNE plot of 2544 successful scRNA-seq libraries from eight donors. Each point represents one cell and points are positioned to retain the pairwise distances as determined by pearson correlation of the 500 most highly expressed genes. Cell identity is indicated by marker gene expression. b) Fraction of cells that express the aging associated gene CDKN2A (p16) in juvenile (0-6 years), young adult (21-22 years) and middle-aged (38-54 years) donors increases with age (P=3.1E-3, n=8, linear regression. Bars are mean +-SEM, n=2-3) c) Transcriptional noise in β-cells is plotted by age group. Whole-transcriptome cell-to-cell variability within cell type is higher in cells from adult donors than in cells from juvenile donors. d) Log2 counts per million (CPM) of cell-atypical glucagon transcript in β cells. Cells from older donors display higher abundance of cell-atypical hormone expression. e) Age and stress genes are strongly associated with transcriptional noise. All genes were tested for association with transcriptional noise (linear rank regression), shown are the top genes by coefficient, with P < 1E-15 (FDR corrected). Line is loess fit +-.995 confidence interval. Dots are running mean, k=10.

The large span of donor ages (≈6 decades), allowed us to assess the effect of organismal aging at the single cell level. Expression levels of known markers of organismal aging such as CDKN2A (p16) were strongly associated with age, but overall we observed only modest systematic age-dependent transcriptional changes for other genes (Fig. 1b, S1a-b). From investigations on a small panel of genes in the mouse heart ^2^, it has previously been suggested that aging is the result of an increase in transcriptional instability rather than a coordinated transcriptional programme. To test whether this observation can be generalized to a full transcriptional profile in human pancreas, we measured the transcriptional noise within cell types and donors using estimates based on Euclidean distance (Fig. S1h) or Pearson correlation as a fraction of technical error (Fig 1c). Both methods indicated increased transcriptional noise in samples from older donors compared to samples from young adults and children, demonstrating age-dependent transcriptional noise.

A subset of α-cells and β-cells simultaneously expressed both *Insulin* (INS) and *Glucagon* (GCG) mRNA - a result which is consistent with prior studies ^6–8^, and which we verified using *in situ* RNA staining (Fig. S1, c-g). scRNA-Seq revealed that the fraction of α- or β-cells co-expressing both *Insulin* and *Glucagon* mRNA increased significantly with advancing age (Fig. 1d, GCG in β-cells: *P*=1.74e-27, n=348. INS in alpha cells [not shown]: *P*=5.38e-10, n=998, linear regression). Thus, increasing numbers of cells with ‘atypical’ hormone mRNA expression is emblematic of age-dependent transcriptional instability, and suggests a physiological basis for declining endocrine function observed by others in aging pancreas ^9,10^.

To investigate whether any systematic gene expression differences accompany an increase in transcriptional noise we performed linear regression on gene expression levels as a function of noise rank (batch corrected and within celltype). As shown in Fig 1e stress response genes such as FOSB, HSPA1A and JUND were most highly associated with increasing transcriptional noise, supporting an aging paradigm that implicates cellular stress in age-related pathology ^11^.

Aging is accompanied by the accumulation of somatic DNA substitutions and the pattern of somatic substitutions in a cell depends on the mutagenic processes that cause them. A growing body of data from tumor genomes has uncovered a multitude of such mutational signatures ^12–15^, many of which can be linked to specific mutational processes. However, these signatures are dominated by processes associated with tumor growth and only three such signatures are linked to aging in tumors or organoid cultures of stem cells ^16,17^. In order to directly study mutational signatures that are active in healthy tissue, we developed a computational method for determining somatic variation within single cells using single-cell RNA-seq data (see methods). Using this method, we compiled a catalogue of putative somatic and germ-line mutations from the 2544 pancreas cells together with 398 previously published single cells from adult human brain ^18^. We also compiled a similar catalogue of clonal variation within 73 GP5d colon adenocarcinoma cells cultured *in vitro*. Germline variation contributed 73.5% of the total number of substitutions on average in pancreas (Fig. 2a). Somatic substitutions were enriched in untranslated regions of transcripts and also enriched for mutations resulting in codons which do not alter the amino acid sequence (Fig 2b, S4h). As expected, the vast majority of putative somatic substitutions were observed in only one cell each (Fig. S2a), indicating that the method is specific to somatic variation. These somatic mutation rates exceed technical error rates due to amplification and sequencing error, as measured by internal spike-in controls of synthetic RNA included in each single cell experiment.

**Figure 2.**
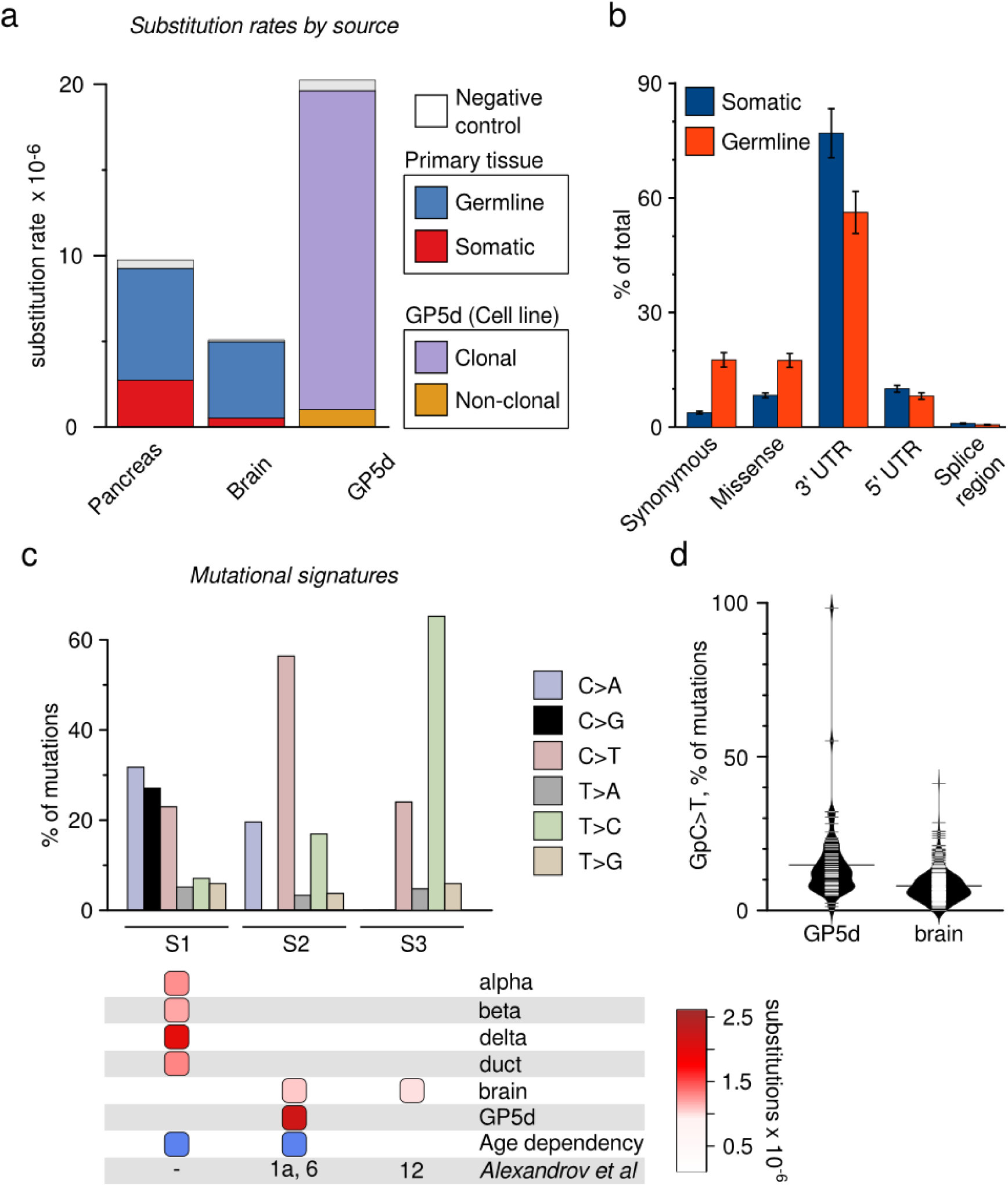
Somatic mutation rates and age-dependent somatic mutational signatures.derived from scRNA-seq data. a) Substitution rates for each type of substitution in the three datasets. Somatic substitution rates are more than five times as high in pancreas as in brain (2.74 × 10^−6^ vs 0.52 × 10^−6^), whereas germline substitution rates were similar between the two. As expected, the rate of clonal substitutions in the tumor cell-line (GP5d) is several fold higher than germline rates in primary tissue. b) Somatic substitutions are strongly enriched on untranslated regions compared to germline substitutions. Bars are mean +- SEM. (3’UTR: P=1.40E-32, paired t-test, n=73) c) Single-nucleotide substitutions in 3003 cells from pancreas, brain and the colon cancer cell line GP5d were organized into five mutational signatures using nonnegative matrix factorization followed by agglomerative hierarchical clustering. Barplot illustrates the percent of mutations attributed to each substitution type in each of the three signatures (S1-S3). Bottom panel denotes the presence of a signature (columns) in a cell type (rows), with color scale indicating strength of signature as median substitution rate for cells of the indicated type. Blue boxes denotes a significant association between signature load and donor age. Bottom row indicates equivalent signatures from ^12^. d) Signature S2 is composed of two sub-signatures corresponding to cancer signatures 1 and 6. Violin plot show C>T substitutions with a preceding G as a fraction of all substitutions in a cell, which is a hallmark of cancer signature 6.

To investigate the patterns of somatic mutations, we determined the rates of the six possible single nucleotide substitutions. Single cells from pancreas had a markedly higher rate (> 5-fold) of somatic variation compared to brain tissue for most substitution types (Fig. 2a, S2c), and there was considerable variation also between cell types in the pancreas (Fig S2b). However, rates of C>T substitutions in a CpG dinucleotide context, known to deaminate spontaneously when methylated, and T>C substitutions were relatively higher in brain compared to pancreas (Fig S2c), in line with what was previously found for postmitotic brain cells^19^. Synthetic control RNA substitution rates were similar between cell types of the pancreas and represent a lower level of technical noise in the measurement. Thus, analyzing the raw sequence reads from scRNA-seq data allows us to determine the mutational history of primary tissues as well as the clonal variation in a tumor cell line.

To identify the mutational signatures (S1-S3, SC4-7) that underlie the observed substitution rates, we used non-negative matrix factorization followed by hierarchical clustering (similar to ^20^, see *methods* for details) on the substitution rates of single cells (Fig. 2c, S3a-c). The S1 signature (high rate of C>A, followed by C>G and C>T substitutions), and S3 signature (highly elevated rate of T>C substitutions), were cell type specific signatures, with S1 found in the endocrine pancreas and S3 in the brain.

The S2 signature was highly enriched in clonal variation within the mismatch repair deficient GP5d cell line, with weaker signal in brain. The pancreas specific signature S1 was characterized by C>A substitutions, with C>G and C>T substitutions at progressively lower rates. C>A and C>G substitutions are attributed to oxidation of the guanine base, creating 8-Oxo-2'-deoxyguanosine (8-Oxo) which mispairs with adenine and can be further oxidized to mispair with guanine ^21,22^, whereas C>T substitutions are attributed to oxidation of the cytosine base^23^. Consistent with oxidation of guanosine driving the mutational signature of β cells, 8-hydroxyguanosine levels were markedly elevated in the DNA of β cells compared to non-islet cells, while only modestly elevated in RNA (Fig. 3). 8-Oxo substitutions preferentially occur when the guanine is on the non-transcribed strand ^24,12^, possibly due to transcription-coupled nuclear excision repair of adducts on the transcribed strand^25^. In order to determine if transcriptional strand bias occurred in our data, we annotated the single-base substitutions with whether the mutated pyrimidine was on the transcribed (-) or untranscribed (+) strand. As expected, C>A and C>G substitutions had a strong preference to occur on the transcribed strand in endocrine cells but not in brain cells, consistent with guanine oxidation driving signature S1 (Fig. S3d). Taken together, signature S1 appears to be a novel, strand-specific mutational signature which bears the hallmarks of oxidative damage.

**Figure 3.**
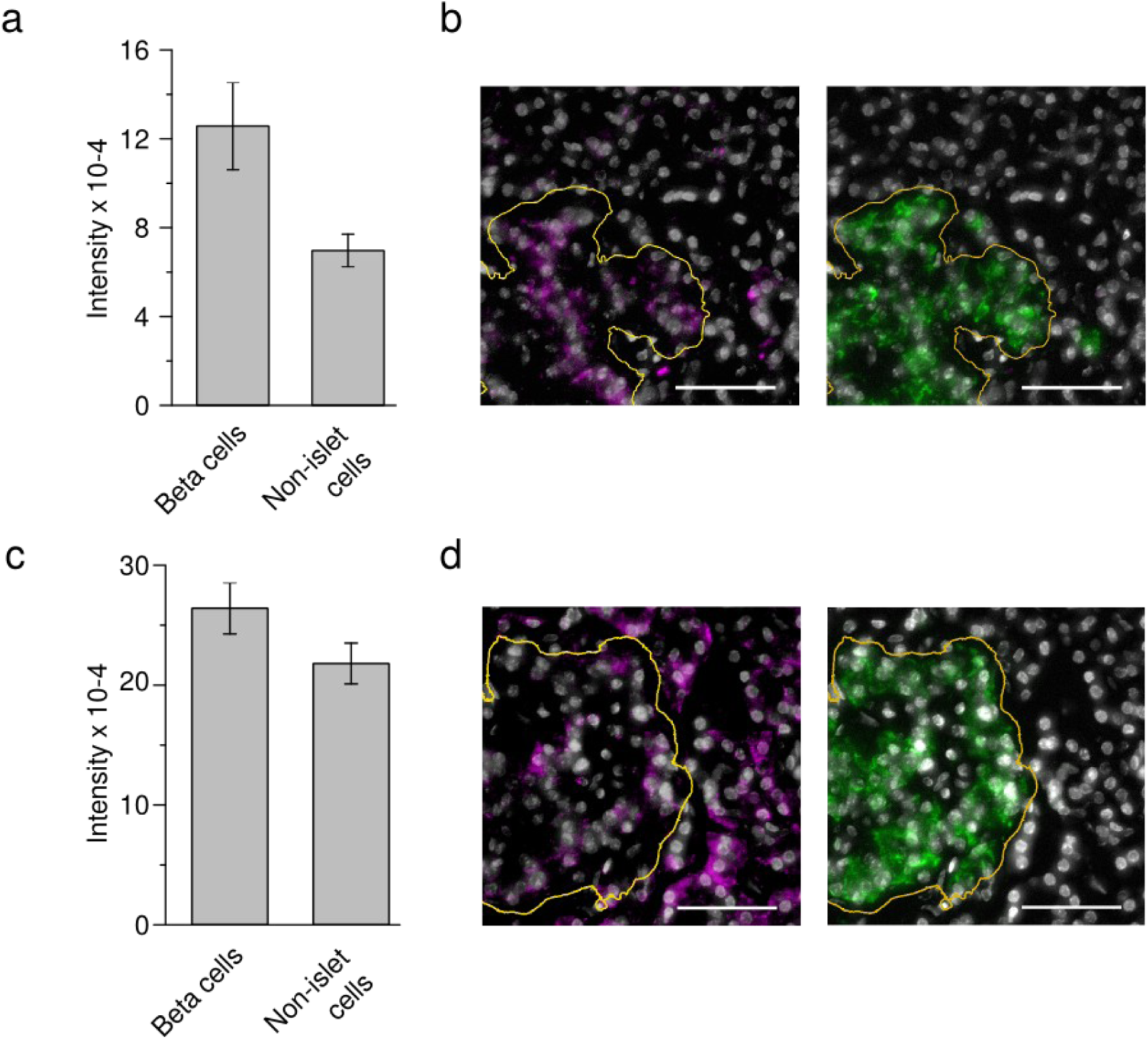
The genome in pancreatic islets are highly enriched in oxidized guanine. a) Pancreatic β-cell DNA is enriched in oxidized guanosine. Nuclear staining intensity of anti 8-Oxoguanosine antibody was quantified for INS-positive β or INS-negative, from the same images. Slides were treated with RNase so as to only measure oxidized bases on DNA. Barplot indicates mean +-SEM ( P=7.30E-57, wilcoxon text. n=769 β-cells, 10713 non-islet cells.). b) Left panel, representative micrograph with 8-Oxoguanosine in magenta and nuclear stain (DAPI) in grey (scale bar, 50μm). Right Panel, Insulin protein staining (green) of the same slide. Insulin-positive islet cell mass is at bottom left, boundary indicated with orange line. c) Pancreatic β-cell RNA is marginally enriched in oxidized guanosine. Cytoplasmic staining intensity of anti 8-Oxoguanosine antibody was quantified for INS-positive β cells and INS-negative cells from the same slides. Barplot indicates mean +-SEM ( P=9.5E-22, 1239 β-cells, 21048 surrounding cells). d) Representative micrograph of 8-Oxoguanosine (magenta, left) and Insulin (green, right). All settings are as in (c,d).

Previous large-scale efforts to decipher cancer-specific mutational signatures in bulk tumor genomes ^12^ discovered 21 unique signatures using the substitution type and the surrounding two bases. We reasoned that our signatures might have been also detected in the tumor data, and compared the signatures by collapsing their probabilities into single-base substitution probabilities. Signature S3 found in this study was very similar to tumor signature 12 from ^12^ (Fig S4d), and the characteristic T>C substitutions in brain display a similar degree of strand specificity to tumor signature 12 (S3d). Signature S2 was almost identical to both the age dependent tumor signature 1, and the mismatch repair associated tumor signature 6 (Fig S4d). The major distinguishing feature between the two tumor signatures is the rate of C>T substitutions within a GpC context, which is higher in tumor signature 6. As shown in figure 2d this distinguishing feature clearly separates the two tissues, suggesting that non-clonal substitutions in GP5d mainly stem from faulty mismatch repair, whereas somatic substitutions in brain are caused by the same age dependent process as tumor signature 1. Interestingly, tumor signature 5, which is of unknown aetiology and is found at low levels in all tumor types is highly reminiscent of our false positive signature (Fig. S4c) - suggesting that it is either a product of false-positive calls in the tumor datasets, or caused by a mechanism that is shared between human replication and enzymes used for nucleic acid amplification. None of the 21 tumor signatures found to date is directly related to endogenous oxidative stress, and the endocrine signature S1 has no direct counterpart among the tumor signatures. Further investigation into tumor signatures of healthy tissues will be needed to elucidate whether signature S1 is emblematic of mainly post mitotic cells with high rate of metabolism, which rarely form tumors, or if it is specific to endocrine pancreatic cells.

Ranking of cells by signature-specific mutational load indicated that signatures S1 and S2 were highly correlated with age, with S1 showing the highest significance (*P*=5.95E-23, Fig. 4a, S4). Only two genes, PON2 (a membrane protein with a putative antioxidant activity) and EGR1, were significantly associated with mutational load of the age dependent S2 (Fig S4a-c). Signature S1, on the other hand, was associated with a large transcriptional effect. The genes most highly associated with high S1 load were involved in transcription (TCEB2), protein synthesis (RPL6) and modulation of ROS (ROMO1) (Fig. 4b). Gene set enrichment analysis indicated that pathways involved in protein synthesis are altered in both cells with high S1 load and high transcriptional noise (Fig 4c). Age-dependent decline in function and regenerative potential has been attributed partially to the activity of reactive oxygen species (ROS) produced by cellular metabolism ^11^. The age-dependent mutational signature in the endocrine pancreas is characterized by a high rate of C>A and C>G substitutions, which are selectively induced by ROS (Fig S4e) ^22,26–27^. Pancreatic islet cells are sensitive to ROS due to low expression of antioxidant enzymes such as SOD1 ^28^, a relatively high rate of ATP dependent processes such as protein production and secretion, and the requirements for reducing power to keep insulin disulfide bonded. Our results thus suggest that the age specific mutational signature observed in the endocrine pancreas is due to ROS dependent lesions on DNA. Interestingly, oxidative damage is part of the pathology of type II diabetes, and plasma 8-hydroxyguanosine is a good correlate to endocrine dysfunction^29^.

**Figure 4.**
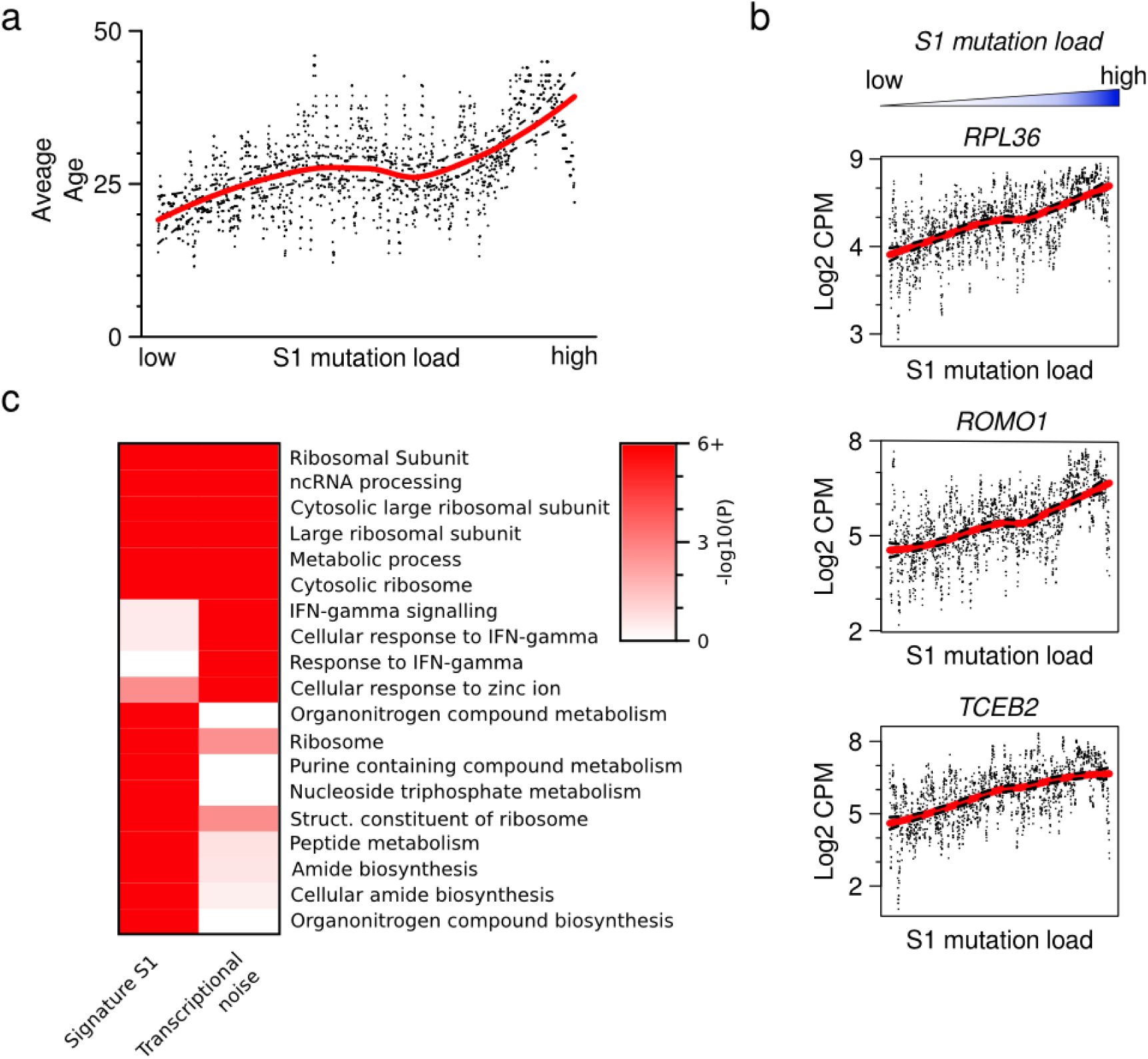
Transcriptional correlates of mutational signatures. Endocrine pancreas cells were ordered according to the fraction of mutations attributed to Signature S1. a) Average age is higher in cells with high S1 load (*P*=5.95E-23, linear rank regression). Points are running mean, k=10, and line is loess fit, dotted lines indicate +− .999 confidence interval. b) Each gene was tested for association with signature S1 (linear rank regression), shown are the top genes by coefficient, with P < 1E-15 (FDR corrected). Points are individual mRNA measurements, line loess fit as in (a). c) Comparison of the top ten gene ontology (GO) categories positively correlated with signature S1 and transcriptional noise. Categories related to protein production, such as ribosomal proteins, recur in both. Color scale indicates FDR-adjusted p-value, winsorized at 10^−6^. The GO term “Ribosomal Subunit” occurred in both lists.

Given that DNA sequence directly influences gene expression, it is tempting to speculate that transcriptional noise in β cells results from an oxidative environment which directly causes lesions to guanine in genomic DNA. However, the correlation between transcriptional noise and signature S1 was weaker than that of either of the two with age, suggesting that they might be caused by independent age related processes. Also, high levels of transcriptional noise seem to occur in younger adults than high levels of oxidative damage (compare figures 1e and 4a). Thus our single cell approach indicates that although the two effects are individually linked to age, DNA damage and transcriptional noise might not be in direct causation - contrary to what has been previously suggested^4^. In a broader sense, our methods for determining transcriptional noise and mutational signatures from scRNA-seq data provide a means to study such signatures in arbitrarily specific cell populations from primary tissue, irrespective of the replicative potential of the cells, which could have far reaching implications on our ability to study such processes.

## Methods

### Human Pancreas and Islet Procurement

All studies involving human pancreas or islets were conducted in accordance with Stanford University Institutional Review Board guidelines. De-identified human pancreata or islets were obtained from previously healthy, non-diabetic organ donors with BMI<30, less than 15 hours of cold ischemia time, and deceased due to acute trauma or anoxia. Organs and islets were procured through Integrated Islet Distribution Network (IIDP), National Diabetes Research Institute (NDRI), UCSF Islet Isolation Core (San Francisco, CA USA) and International Institute for the Advancement of Medicine (IIAM). For FACS, scRNA-Seq studies islets from three juvenile (ages 1 month-old, 5, 6), and five adult donors (ages 21, 22, 38, 44, 54 years) were used. For immunostaining studies pancreatic tissue sections from a 31-year-old donor were used.

### Flow Cytometry

Isolated human islets were dissociated into single cells by enzymatic digestion using Accumax (Invitrogen). Prior to antibody staining, cells were incubated with blocking solution containing FACS buffer (2% v/v fetal bovine serum in PBS and goat IgG [Jackson Labs], 11.2 μg per million cells). LIVE/DEAD Fixable Aqua Dead Cell Dye (Life Technologies) was used as a viability marker. Cells were then stained with appropriate antibodies at 1:100 (v/v) final concentration. The following antibodies were used for FACS experiments: HPx1-Dylight 488 (Novus, NBP1-18951G), HPi2-Dylight 650 (Novus, NBP1-18946C), CD133/1 - Biotin (Miltenyi Biotec 130-090-664), CD133/2 - Biotin (Miltenyi Biotec 130-090-852), streptavidin-eFluor780 (eBioscience, 47-4317-82), streptavidin-APC (eBioscience, 17-4317-82), anti human EpCAM-APC (Biolegend, 324208). Cells were sorted on a special order 5-laser FACS Aria II (BD Biosciences) using a 100 m nozzle following doublet removal. Sorted single cells were collected directly into 96-well plates (Bio-Rad cat #: HSP9601) containing 4 µL of lysis buffer with dNTPs ^5^ for downstream single-cell RNA-Seq assays.

### Single-Cell RNA-Seq and Data Analysis

Single-cell RNA-Seq libraries were generated as described ^5^. Briefly, single-cells collected in 96-well plates were lysed, followed by reverse transcription with template-switch using an LNA-modified template switch oligo to generate cDNA. After 21 cycles of pre-amplification, DNA was purified and analyzed on an automated Fragment Analyzer (Advanced Analytical). Each cell’s cDNA fragment profile was individually inspected and only wells with successful amplification products (concentration higher than 0.06 ng/ul) and with no detectable RNA degradation were selected for final library preparation. Tagmentation assays and barcoded sequencing libraries were prepared using Nextera XT kit (Illumina) according to the manufacturer’s instructions. Barcoded libraries were pooled and subjected to 75 bp paired-end sequencing on the Illumina NextSeq instrument.

Sequencing reads were trimmed, adapter sequences removed and the reads aligned to the hg19 reference assembly using STAR ^30^ with default parameters. Duplicate reads were removed using picard ^31^. Transcript counts were obtained using HT-Seq ^32^ and hg19 UCSC exon/transcript annotations. Transcript counts were normalized into log transformed counts per million (log2(counts * 1 000 000 / total_counts + 1). Single cell profiles with the following features were deemed to be of poor quality and removed: 1) cells with less than 100.000 total number of valid counts on exonic regions. 2) cells with very low actin CPM. To determine a cutoff for actin CPM, we used the normal distribution with empirical mean and standard deviation from actin. The cutoff was set to the 0.01 quantile (eg. the lower 0.01 % of the bell curve).

Pairwise distances between cells were estimated using Pearson correlation on the 500 most highly expressed genes in any one cell. Dimensionality reduction of the pairwise correlation matrix was performed using the t-SNE method ^33^.

To determine Gene Ontology categories that were associated with transcriptional noise or signature specific mutational load, we used Gene Set Enrichment Analysis (GSEA), using the coefficients of association to noise/rank of significantly altered genes (P<1E-5, linear model, FDR corrected). Coefficients were used as a preranked list in the GSEA software using default parameters with the gene set database “c5.all.v5.2.symbols.gmt”, which includes all GO categories.

The data reported in this paper have been deposited in the Gene Expression Omnibus (GEO) database, accession no. GSE81547.

### Genomic sequencing

Genomic variants were determined from whole genome sequencing data following GATK Best Practices^34^. Briefly, adapters and low quality bases were trimmed using cutadapt v1.9^34,35^. Reads were aligned to hg19 using BWA-MEM 0.7.12 ^36^. Duplicates were removed using Picard tools v1.119 followed by indel realignment and base recalibration using GATK v3.5^31^. Variants were called using haplotype caller and recalibrated using VQSR. Default software parameters were used and reference files downloaded from the GATK Resource Bundle 2.8/hg19.

### Somatic mutational signatures in single-cell RNA-seq data

To explore mutational signatures in single postmitotic cells, we analyzed the raw sequence reads from mRNA-seq. Previously, mutational signatures have been successfully extracted from exon sequencing; however using single-cell data poses a number of additional challenges. First, we need to deal with the higher error rate associated with reverse transcription and a higher number of PCR cycles. We do this in two ways - by including positive and negative internal controls for each cell, that are used to derive a meaningful cutoff when calling substitutions, and by performing an additional post-selection of signatures, discarding potential false-positives. Second, the sequence space in a single-cell RNA-seq experiment is typically fairly limited, even compared to exon sequencing. We mitigate this issue by sequencing long reads (75 bp paired-end), and by sequencing deeper than typically needed for scRNA-seq (approx. 1M mapped reads per cell). Further, we calculate substitution rates based on the actual number of sequenced kmers in each cell, to account for differences in base distribution. Finally, the limited number of substitutions in each cell means that the sequence context cannot be reliably included in all cases, which is why we generally restricted ourselves to analyzing single-base substitutions.

Raw variation calls were made using the Haplotype Caller (GATK pipeline^31,34^) on the BAM files after applying SplitNCigarReads to remove overhangs into intronic regions. Variants were filtered to remove clusters (>3 SNPs within 35 bases), as well as variants with QD < 2.0 and FS > 30.0. Germline mutations were called using a merged set of all single-cell profiles from each patient. Subsequently, we filtered the raw variation calls by applying variant quality score recalibration using the GATK pipeline. To reliably call substitutions we need internal controls for each cell, corresponding to a true-positive and true-negative set. We used known variants (dbSNP release 138) from our germline calls that mapped to transcribed regions of the genome as a true positive set (phred-scaled prior: 15.0) and variants that map to ERCC control reads as a false positive set (ERCC controls are synthetic RNA sequences and therefore devoid of systematic variation). To filter somatic substitutions, a strict cutoff, allowing 10% false negative rate was used. Variants also found in the germline were flagged as germline mutations and not used for somatic signatures. In all subsequent analysis, only single-nucleotide substitutions were considered.

For each cell, we extracted the genomic context of each mutation and created a catalogue of the frequency of mutation types. We then divided these frequencies with the kmer counts derived from fastq sequences for the cell to obtain the final substitution rates. Negative control ERCC sequences were processed in parallel, to give accurate substitution rates that reflect the different sequence background. Substitution rates in these ERCC samples were 4.8E-7. If we assume that the false-positive substitutions stem exclusively from somatic calls (eg. that the germline calls are completely devoid of false positives), this result would indicate a false discovery rate of 15.05% for somatic substitutions. Thus, we estimate that the upper bound of our false discovery rate is 15%. To further validate our method we performed 25x whole genome sequencing (WGS) of GP5d and compared the overlapping substitution calls from single-cell mRNA seq and bulk genomic sequencing. A total of 151,030 genomic positions were determined to have single-base substitutions from the reference genome based on mRNA-seq. Out of these 151,030 substitution calls, 105,673 were also found in WGS and 105,543 were identical (concordant). 45 357 substitutions, or 30.0% of total, were not found in WGS calls; these calls include somatic substitutions, false negative calls from WGS and technical errors. Based on these numbers and our findings about somatic substitution rates, we estimate that the false positive rates due to technical errors are accurately described by the ERCC substitution rates.

To determine how well our method identifies somatic substitutions, we again used bulk WGS as a gold standard. We determined the overlap of WGS substitution calls with all putative somatic substitutions called from the merged mRNA-seq data. Out of 4637 putative somatic substitutions 612 were also found in WGS, indicating that 13.2% of our putative somatic calls are actually germline SNPs.

Thus, we estimate the overall false discovery rate in our data (before applying nonnegative matrix factorization and signature selection) to be approximately 25%, which includes 13.2% that represent real variation stemming from germline rather than somatic events and ~10-15% substitution calls that were erroneously called due to technical errors such as PCR or sequencing artifacts.

To further explore structure within the somatic substitution calls, we examined the effect of substitutions on protein sequence. Because of the degeneracy of the exon code, a fraction of exonic substitutions will give rise to a DNA sequence which codes for the same amino acid sequence. Such synonymous (or silent) substitutions are enriched in germline SNPs, and given that a subset of amino acid substitutions will negatively affect fitness of the cells, we would expect some enrichment of synonymous substitutions also among somatic substitutions. Also, we would expect this enrichment to be similar in different cell types, irrespective of the mutational load. Substitution calls due to technical errors, however, will not be enriched in silent substitutions. We annotated the substitution calls based on genomic notation (hg19), and calculated the fraction of calls that result in a codon for the same amino acid. As a comparison, we calculated the fraction of synonymous substitutions based on random DNA mutation. The average fraction of synonymous substitutions was 40% higher than expected by random chance (0.32 in pancreas compared to 0.23 expected by random, P=3.34E-125, Wilcoxon test. Figure S4h). Importantly, this number did not correlate with mutational load; cells with higher number of mutations in fact had a somewhat increased fraction of synonymous substitutions (Slope=3.25E-5, P=0.08, linear regression), and pancreas cells had almost identical fraction of silent mutations compared to brain even though the substitution rate was five-fold higher in pancreas (Fig S4i). Thus, the differences in substitution rates likely reflect genetic alterations in the cells, rather than technical error.

To decipher the underlying mutational signatures, we applied non-negative matrix factorization using the NMF R package ^37^ to the substitution rates of single-nucleotide substitutions (eg. the mean of the rates for a substitution type over all contexts) for each cell type separately. The highest scoring solution out of 10000 independent runs of the algorithm was used for the final result. The number of possible signatures (5) was chosen to be higher than the number of unique signatures actually found by the algorithm, and duplicate signatures were merged together. We applied hierarchical clustering on the full set of mutational signatures (“basis matrices”) to identify distinct mutational signatures (Fig S3a). Finally, we selected signatures based on five criteria (summarized below and in Fig S3c). To find the signatures that likely represent cell type specific processes that were active in the healthy cell during the donor’s lifetime, we determined cell type specificity and age dependence of each signature. Also, because of the relatively high level of noise in the data, a signature might represent errors that arose systematically during reverse transcription. Thus, to arrive at the final three signatures (S1-S3), removed mutational signatures with a high degree of similarity to the substitution rates of the negative control RNA, with no cell-type specificity, positive age dependence, or with a very low signal. We also determined the similarity of the signatures to the COSMIC tumor signatures^12^. Figure S3C, bottom panel, summarizes the association of signatures with these traits. It should be noted that we cannot completely rule out the possibility that the excluded signatures were due to a cell-type specific process active during the lifetime of the donor. Further investigation on much larger panels of tissues will be needed to determine the origin of these signatures.

Fig. 2b and S3c show the geometric median signature of each cluster. Mutational load of a signature on a cell was determined as the fraction of somatic substitutions of that cell attributed to the signature in question. To obtain a signature load ranking, cells were ordered according to the fraction of mutations that are attributed to a specific signature. Statistical significant association was determined using linear regression.

### Estimation of transcriptional noise

In order to ascertain the robustness of age dependent transcriptional noise, we computed three measurements of transcriptional instability each of which displayed a strong statistical significance and positive coefficient to age. For per-donor measurements we first divided the cells into cell types and computed the mean expression vector for each cell type. We then calculated the Euclidean distance between each cell and its corresponding celltype mean vector. The individual datapoints are summarized as boxplots. As an alternative method to obtain a measure of the transcriptional noise of a single cell, we first subsampled the gene count list to 100 000 counts per cell. We then selected a set of invariant genes evenly across the range of mean expression. First we binned the genes in 10 equally sized bins by mean abundance, then we selected the 10% of genes with the lowest CV from each bin, omitting the bins at the high and low extremes. We then used these genes to determine the Euclidean distance from each cell to the average profile across all cells. Finally, we used a correlation based method where noise is expressed as biological variation over technical variation. First, we calculated the biological variation b_ijk_=1-cor(x_ijk_, u_ij_), where ui is the mean expression vector in cell type i, patient j and x_ijk_ is the expression vector of cell k in that cell type i, patient j. Next, we calculated the corresponding technical variation t_ijk_=1-cor(x^contr^_ijk_, u^contr^) where x^contr^_ijk_ and u^contr^ are the expression vector and mean expression vector of the ERCC spike-in controls. The final measurement is b_ijk_/t_ijk_ eg. the biological noise as a fraction of technical noise. The cells were ordered by this distance within cell type, and their ranking used for linear regression.

To determine the genes whose mRNA abundance were significantly dependent upon transcriptional instability, we used linear rank regression on the CPM values. P-values were adjusted for multiple testing using the FDR procedure of Benjamini & Hochberg (with FDR < 1E-15 as significance cutoff), and ordered by their coefficient.

### *In situ* RNA and protein staining

Multiplex RNA staining was performed on 10 µm thick, formalin-fixed, tissue sections using barcoded transcript-specific padlock probes and rolling circle amplification (RCA) as described before ^38^. The primer sequences were

**GCG:** G+TC+TC+TC+AA+AT+TC+ATCGTGACGTTT

**INS:** G+CA+CC+AG+GGC+CCC+CGCCCAGCTCCA

Padlock probes:

**GCG:** Phosp-GAATAACATTGCCAAACGTGTGTCTATTTAGTGGATCCCGTGCGCCTGGTAGCAATTAGCTCCACTGTTACTAGATTGGAATACCAAGAGGAACAG

**INS:** Phosp-AGGTGGGGCAGGTGGAGCCTCAATGCTGCTGCTGTACTCTACG

ATTTTACCAGTTGCCCTAGATGTTCCGCTATTGTCCGGGAGGCAGAGGACCTGC

Detection probes:

**DO_1_FITC:** AGUCGGAAGUACTACTCUCT_FITC

**DO_1_Cy3:** CCUCAATGCUGCTGCTGUAC_Cy3

**DO_1_Cy5:** TGUGTCTATUTAGTGGAUCC_Cy5

**DO_2_FITC:** CGUGCGCCUGGTAGCAAUTA_FITC

**DO_2_Cy3:** AGUAGCCGUGACTATCGUCT_Cy3

**DO_2_Cy5:** TCUACGATUTTACCAGTUGC_Cy5

**DO_3_FITC:** CCUAGATGTUCCGCTATUGT_FITC

**DO_3_Cy3:** GCUCCACTGUTACTAGAUTG_Cy3

**DO_3_Cy5:** CTUGTGCTGUATGATCGUCC_Cy5

and accession numbers of the probes used are reported in Supplementary Table 1 The RCA products were stained by sequential hybridization of three uracil-containing fluorescent oligonucleotides following a modified protocol from Ke 2013. The three reported probes were mixed 0.1 mM each with hybridization buffer (20% formamide in 2x SSC) and incubated with the tissue at 37°C for 30’. After incubation, tissue section was washed in PBS 5’ and nuclei were counterstained with DAPI 300nM in PBS at room temperature for 15’. The tissue was washed in ethanol 70, 85 and 100% 5’ each, air dried and mounted in Antifade gold (Invitrogen) before imaging. After imaging, the fluorescent probes were removed by digestion with 0.02 U/µl UNG (Thermo) in UNG buffer and 0.2µg/µl BSA at 37°C for 30’ followed by two washes in 65% formamide pre-warmed at 55°C. Consecutive staining of the RCA products were performed, in the same way, with different set of fluorescent probes.

**Table 1.** Summary of sequenced cells, only cells with > 10E5 aligned counts are shown. Sequencing statistics are median values.

|  | Passed QC | Failed QC |
| --- | --- | --- |
| Cells | 2544 (94.9%) | 136 (5.1%) |
| Sequencing statistics |  |  |
| aligned reads | 932172 | 962153 |
| transcripts detected | 3203 | 1392 |
| % aligned | 78.54% | 79.94% |
| % ERCC | 8.06% | 33.20% |
| % exonic (non-ERCC) | 62.85% | 29.03% |
| % mitochondrial | 6.47% | 10.53% |

After RNA, immunofluorescent staining was done on the same tissue section. The tissue was washed twice in PBS with 0.025% triton X-100 at room temperature and blocked with 1%BSA in PBS for 2 h at room temperature. Antibodies against human Insulin (DAKO, A0564, guinea pig) and glucagon (Sigma, G 2654, mouse) were diluted 1% in PBS containing 1% BSA and applied to the tissue and incubated at 4°C overnight. The tissue was washed twice in PBS with 0.025% triton X-100 before incubation with 1% anti-guinea pig GFP labelled and anti-mouse Cy5 secondary antibody, 1% BSA in hybridization buffer for 1 h at room temperature. Cy3-labelled RCA reporter probes were also added at 0.1 µM concentration to stain all the RCA products and used to align immunofluorescence images to previous RNA staining. After incubation in secondary antibody the section was washed 3 times in 1xPBS at room temperature before mounting in Antifade gold and imaging. For 8-hydroxyguanosine staining, 8-oxo-dG Ab (MyBioSource, MBS606843, mouse) was used, which binds to the oxidized based both in DNA and RNA. To measure the levels of oxidized genomic guanine, cells were treated with RNaseA before staining according to the protocol provided by the manufacturer. Briefly, sections were incubated in PBS buffer containing 500 µg/ml RNaseA (ThermoFisher), 150 mM NaCl and 15 mM sodium citrate for 1 h at 37˚C. After washing the sample twice in PBS the DNA was denatured by incubating with HCl 2N for 5’ at room temperature and then neutralized by incubation with Tris-base 5’ at room temperature followed by two washes in PBS. Blocking and antibody staining against human insulin and 8-Hydroxy-2’-deoxyguanosine was performed as described before (anti 8-oxo-dG was used at 1:250 dilution).

Multidimensional imaging was done with a Zeiss Axioplan epifluorescence microscope equipped with filter-cubes for DAPI, FITC, Cy3 and Cy5, a Axiocam 506 mono camera (Zeiss), automated filter-cube wheel and a motorized stage. Z-stacks of 15 images were acquired with a Plan-Apochromat 63x objective and check objective) several field of view of each region of interest were projected (maximum intensity projection) and automatically stitched using the Axiovision software (Zeiss).

Images were exported as single-channel 16-bit grayscale and analyzed as described before ^38^. Briefly, single channels images from staining cycle one were combined and used as mask to align images from subsequent cycles based on nuclei and RCA staining. Image alignment was done using MultiStackReg module of ImageJ (version 1.50e). Pre-aligned RNA images were analyzed with CellProfiler 2.1.1 (rev 6c2d896) and intensity and position of RCA products were measured using the same pipeline as in^39^. The barcode decoding was obtained using the same Matlab script as described before^47^. Lowering the quality threshold to zero (Qt=0) allowed us to increase sensitivity of detection while the fraction of insulin and glucagon signals detected outside the islets (false positives) was still negligible (less than 0.3% of all GCG and INS signals). Object-based measurement of immunostaining intensity was done with CellProfiler on the corresponding images using the identified RCA products as mask.

## Author contributions

M.E., H.E.A, S.K.K. and S.R.Q. designed research; M.E., H.E.A, JB and M.M performed research; R.B. isolated islets; M.E., and S.R.Q. analyzed data; M.E, H.E.A, M.M, S.K.K., and S.R.Q. wrote the paper.

## Acknowledgements

The authors thank Norma Neff and Gary Mantalas for assistance with sequencing and Spyros Darmanis, Geoff Stanley and Felix Horns for helpful discussions. This study was supported by California Institute for Regenerative Medicine Grant GC1R-06673, Center of Excellence for Stem Cell Genomics and National Institutes of Health Grants U01-HL099999 and U01-HL099995 (to S.R.Q.). M.E was supported by the Wallenberg Research link at Stanford University. H.E.A. was supported by a postdoctoral fellowship from the JDRF and a training grant to the Endocrinology Division, Department of Medicine, Stanford (5T32DK007217-39, NIDDK). Marco Mignardi was supported by the Swedish Research Council, grant number XXX-2015-00599.

---

**Figure S1.**
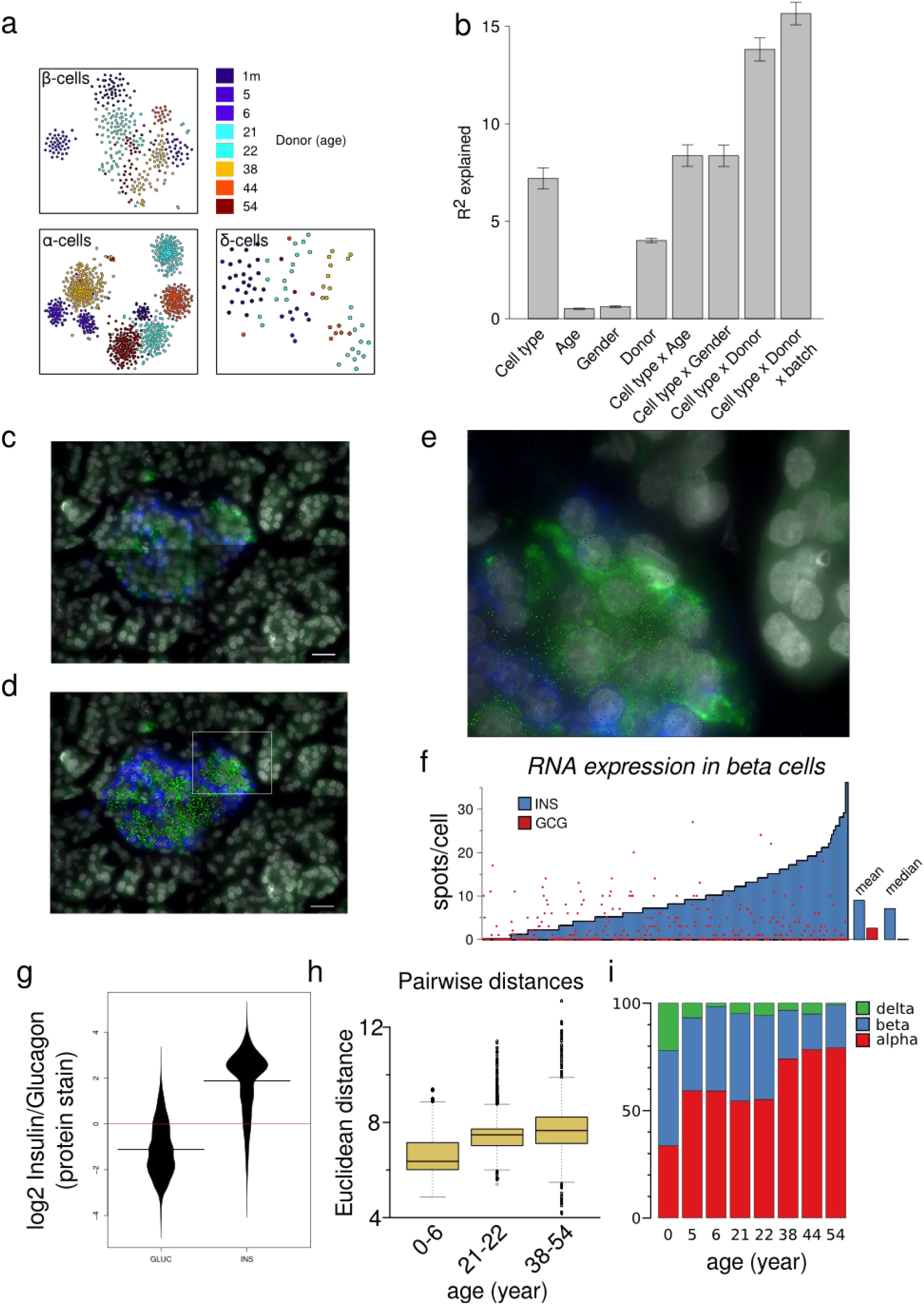
a) tSNE plot of cells from the major endocrine cell types. Colors are by donor (as specified by age, top right panel). Cells cluster by donor suggesting that our data could not find support for sub cell types that have a stronger cell identity than individual variation. b) Relative contributions of cell type, age, gender, donor and library preparation batch. Error bars are mean +/-SEM. c-e) Parallel protein and RNA staining *in situ.* A representative image at 63x magnification of a pancreatic islet containing cells with atypical hormone expression. Scale bar is 20 µm. c) protein stain only (green: insulin, blue: glucagon), d) *in situ* RNA-staining (dots) + protein stain (green dots: INS gene specific, blue dots: GCG gene specific). e) Magnified version of c). f) Quantification of cell-atypical hormone expression *in situ*. Green bars show number of INS spots per cell, red dots number of GCG spots. There was no significant dependency between INS expression and GCG expression (*P*=0.859, linear regression). g) Violin plots of the ratio of Insulin to Glucagon protein staining at the sites of Insulin (INS, *n*=5801) and Glucagon (GCG, *n*=3254) RNA hybridization spots. h) Pairwise Euclidean distances between 10000 random pairs of endocrine cells from each donor is plotted by age group. Whole-transcriptome cell-to-cell variability between β-cells from adult donors is higher than variability between cells from juvenile donors. i) Fractions of endocrine cell types in donors. Donors are sorted by age.

**Figure S2.**
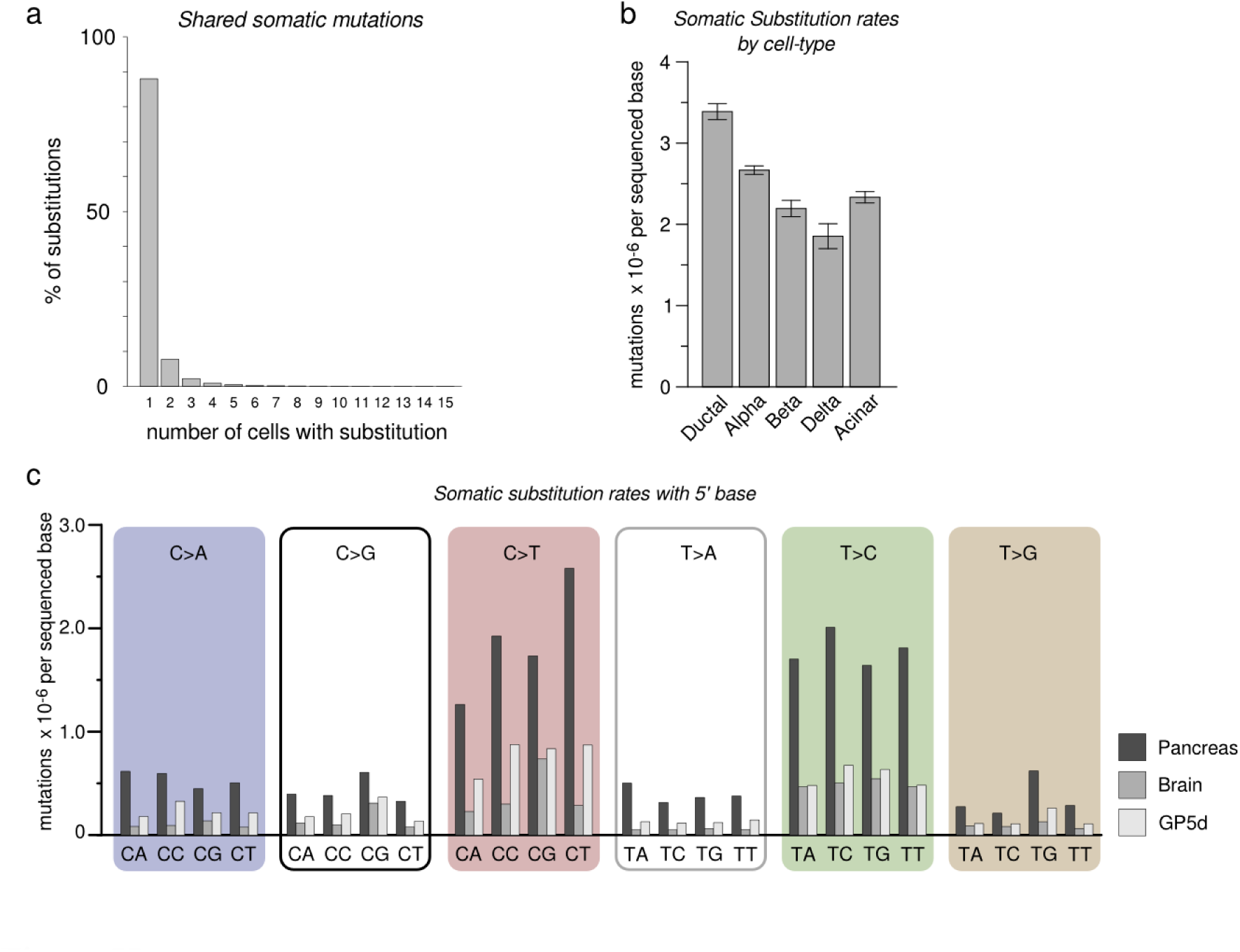
a)The distribution of the number of occurrences of distinct somatic (non-germline) substitutions. As expected, somatic mutations that are shared between more than one cell are rare. b) Somatic substitution rates vary between cell types in the same organ (bars are mean +- SEM). c) Comparison of mutation rates of single-nucleotide substitutions in the context of the nucleotide immediately 5’. Different substitution types are separated by boxes with the substitution type indicated (eg. C>A: C to A transversion). Brain cells display a lower substitution rate overall, although the difference is smaller for C>T substitutions within a CpG context, associated with DNA methylation.

**Figure S3.**
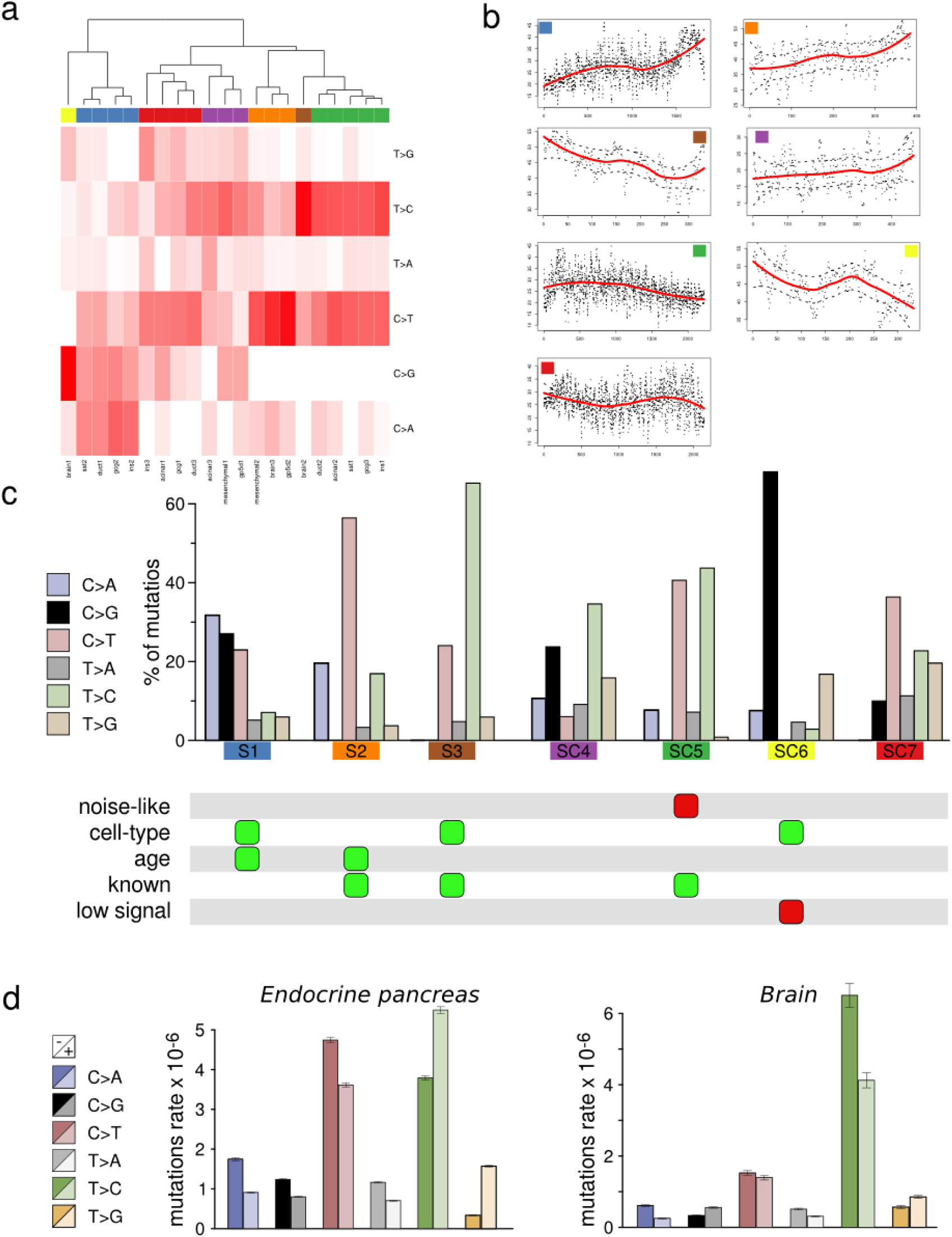
Mutational signatures. a) Heatmap showing raw signatures from non-negative matrix factorization. Dendrogram (top) indicates hierarchical clustering, and clusters at the 6th branch point shown as colored bar between dendrogram and heatmap. The spatial median of each cluster is shown in Fig. 2a and S3c. b) Association of signatures S1-3, SC4-7 to age. Cells were ordered according to the fraction of mutations attributed to the indicated signature. Dots are running mean of age, k=10. Line is loess fit, dotted lines indicate +-.999 confidence interval. c) Fractional barplots of all signatures (S1-3, SC4-7). Colors as in a. Bottom panel indicates selection items for determining whether to exclude the signature. Green: cause for inclusion, Red: cause for exclusion. d) Strand specificity differs between cell types. Mutations were annotated based on if the mutated pyrimidine occurred on the transcribed (-) or untranscribed (+) strand. Bars represent raw substitution counts in endocrine cells (left) and brain cells (right). Note that endocrine cells have a strong strand bias for the transcribed strand for C>A, C>G, C>T substitutions (also found in oxidative stress-related tumor signatures^12^), while brain has a similar bias for the transcribed strand for T>C substitutions (simlilar to tumor signature 12^12^).

**Figure S4.**
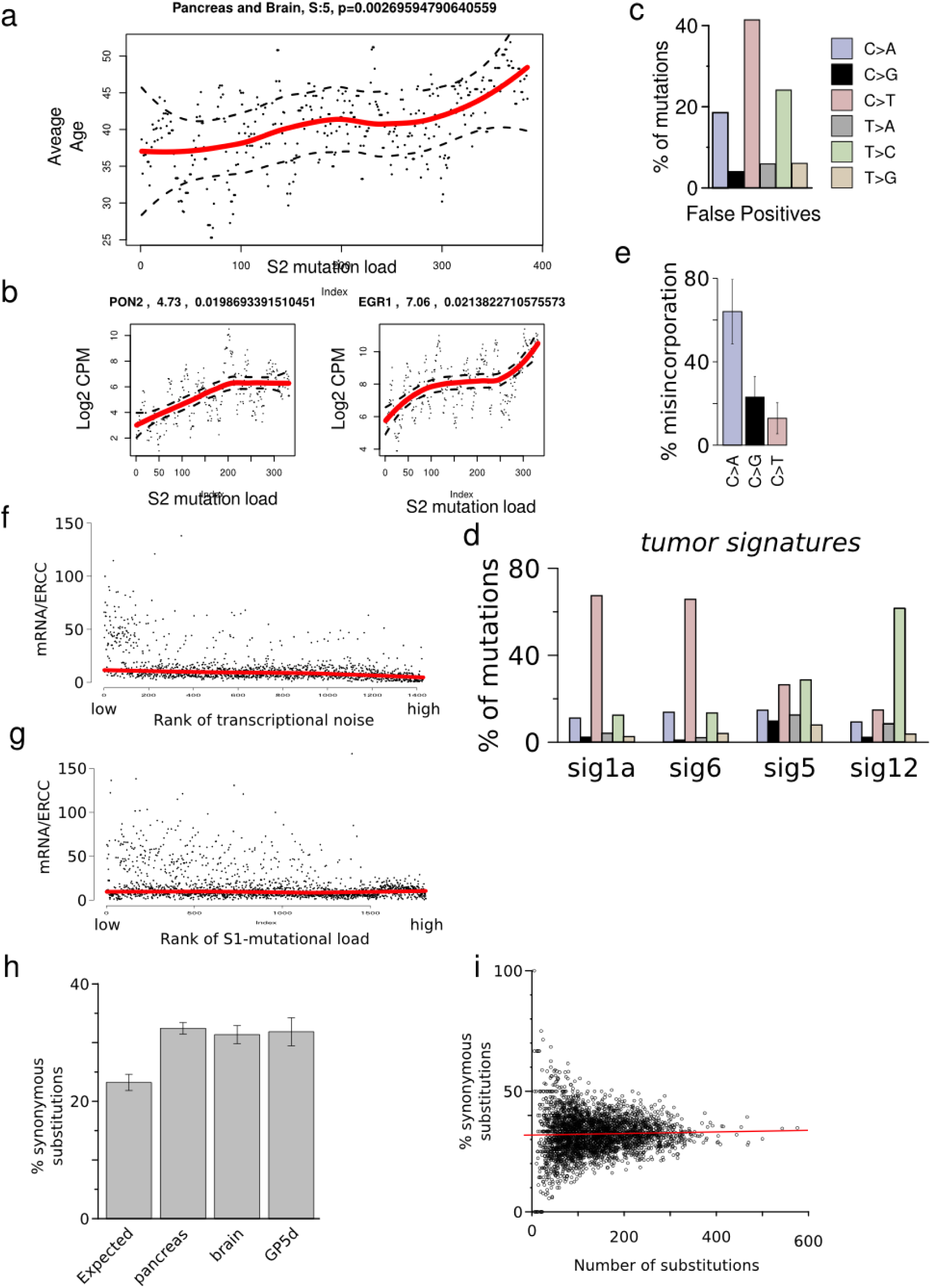
Transcriptional correlates of mutational signatures. Brain cells were ordered according to the fraction of mutations attributed to Signature S2. a) Average age is higher in cells with high signature S2 load (*P*=2.7E-3, n=398. linear rank regression). Line is loess fit +-.999 confidence interval. Dots are running mean, k=10. b) Each gene was tested for association with signature S2 (linear rank regression), shown are the top genes by coefficient, with P < 5E-2 (FDR corrected). Line is loess fit +-.999 confidence interval. Dots are individual observations. c) Signature of raw substitution rates in ERCC spike-in RNA constitutes a false-positive signature. d) Tumor signatures from Alexandrov. et al.^12^, collapsed into substitution types without 3’/5’ context by addition. e) Empirical misincorporation rates caused by 8-Hydroxyguanosine in vitro. Data from from Kamiya et al.^40^. f-g) Ratio of human mRNA to spike in control in cells, ordered by rank of transcriptional noise (f) or rank of signature S1 mutational load (g). h) Synonymous substitutions generating an identical codon as the reference sequence are enriched in somatic variation from all tissues. i) The fraction of synonymous substitutions is not positively correlated with overall mutation load.

